# Yield from the shadows: beyond top layer photosynthesis to enhance crop productivity

**DOI:** 10.1101/2025.08.04.668377

**Authors:** Carlos A. Robles-Zazueta, Gemma Molero, Matthew P. Reynolds, Erik H. Murchie

**Affiliations:** Division of Plant and Crop Sciences, School of Biosciences, University of Nottingham, Sutton Bonington Campus, Leicestershire, LE12 5RD, United Kingdom; Global Wheat Program, International Maize and Wheat Improvement Center (CIMMYT), carretera Mexico-Veracruz km 45, Texcoco 56237, Estado de Mexico, Mexico; KWS Momont Recherche, 7 rue de Martinval, 59246 Mons-en-Pevele, France

**Author notes:** Department of Plant Breeding, Hochschule Geisenheim University, 65366 Geisenheim, Germany (Carlos A. Robles-Zazueta).

**Keywords:** canopy photosynthesis, crop productivity, food security, wheat improvement, yield

## Abstract

Photosynthesis research in crops typically focuses on upper canopy layers, which is partly for convenience and partly for the sake of achieving stable datasets under high light conditions. This neglects significant contributions from light - limited portions of the canopy within the lower layers. This study aimed to provide an empirical quantification of the role of these hidden layers of wheat canopies in the context of canopy scale productivity. We demonstrate that light-saturated photosynthetic rates (A_sat_) in middle and bottom layers at key growth stages can be strong predictors of grain yield. Despite variability in architecture across layers, light interception remained similar and key associations between biomass accumulation and yield with A_sat_ emerged. Yield showed positive associations with photosynthesis in all canopy layers but was stronger at the top layer during grain filling and at the bottom layer during booting. Whole canopy photosynthetic rates were influenced by top layer architecture, N availability in the middle and bottom layers and leaf angles at the bottom of the canopy. Our findings suggest that measurements within hidden layers are required, and that optimizing middle and bottom layer A_sat_ during the vegetative period and top layer A_sat_ during grain filling can boost food security.

## Introduction

Wheat (*Triticum aestivum* L.) is the most widely grown crop worldwide accounting for ∼30% of all sown area for cereals and ∼ 40% of total cereal exports at 118 million tonnes (FAO, 2021). Sustaining and increasing wheat yield is jeopardized by shifts in climate patterns (Langridge & Reynolds, 2021) and socio-economic disruptions such as COVID-19 pandemic and the Russia-Ukraine conflict (Bentley *et al*., 2022). The challenge of increasing staple crop yields, without increasing cultivation area, coupled with the threats posed by climate change represent one of the toughest problems’ humanity will face this century.

From a physiological point of view, crop yield is the product of the photosynthesis of leaves across all layers of the canopy, spikes and stems with each organ being exposed to distinct micro-environmental conditions, which makes photosynthesis associations to yield one of the most complex biological processes to study. Agronomically speaking, yield can be defined as the product of radiation intercepted throughout the crop cycle, the efficiency of its conversion biomass (i.e. radiation use efficiency, RUE) and the proportion of biomass stored in the grains at physiological maturity (i.e. harvest index, HI) under yield potential conditions (Slafer, 2003; Murchie *et al*., 2009; Reynolds *et al*., 2012). Recent work suggests that photosynthesis from lower layers of the canopy could have a large role in improving biomass accumulation and yield in cereals (Burgess *et al*., 2018; Foo *et al*., 2020; Salter *et al*., 2020) and canopy photosynthesis models partially consider this (e.g. WIMOVAC, APSIM, DcAPST; (Casadebaig *et al*., 2016; Song *et al*., 2017; Wu *et al*., 2019)). However, despite the fact that these hidden parts make up a substantial proportion of canopy leaf area, our empirical understanding of their function beyond canopy photosynthesis models is very limited.

A large number of reviews have asserted that increasing photosynthesis will be one of the main pathways to increase crop yield, including wheat, because leaf photosynthesis is the primary source for carbohydrate accumulation in plants for most part of the growth cycle (Horton, 2000; Long *et al*., 2004; Murchie *et al*., 2009, 2018; Zhu *et al*., 2010; Lawson *et al*., 2012; Reynolds *et al*., 2012; Faralli & Lawson, 2019; Evans & Lawson, 2020; Furbank *et al*., 2020; Simkin *et al*., 2020; Tosens *et al*., 2025). However, correlation between measurements of leaf photosynthesis and grain yield has been found to be inconsistent across crops (Supplementary Table 3). This lack of consistency could be partly explained because most studies relating photosynthesis with source or sink traits have focused on short term measurements of light saturated photosynthetic rates in the sunlit layer of the canopy, notably the flag leaf in cereals, which is commonly considered the main active photosynthetic organ contributing to grain yield. In wheat, measuring just flag leaves has been previously justified due to associations found with yield in elite germplasm (Fischer *et al*., 1998). However, this may not allow us to exploit the full phenotypic capability of photosynthesis within currently available germplasm (Murchie *et al*., 2018). Further the emergence of high-throughput methods as proxies for photosynthesis research (Serbin *et al*., 2012) coupled with advanced statistics, photosynthesis and growth analysis presents an opportunity for understanding the functional contribution of canopy layers. To date no study has tried to dissect the complex relationship between canopy layer photosynthesis and yield in the field.

Light availability declines from the top to the bottom of the canopy exponentially which is usually aligned with chlorophyll and N distribution according to canopy optimisation theory (Hirose, 2005; Hikosaka, 2016), whereby the average light intensity should be tuned to the photosynthetic capacity by light acclimation (Murchie *et al*., 2002). Indeed, it has been proposed that optimising canopy photosynthesis could take place by redistributing N toward upper leaves to enhance photosynthetic capacity whilst minimising reductions in the lower leaves (Salter *et al*., 2020) or by reducing antenna size and chlorophyll b content to improve canopy photosynthetic efficiency (Ort *et al*., 2011). Lower canopy layers experience contrasting light conditions compared to upper layers. They are typically shaded and receive low levels of photosynthetically active radiation (PAR) near the light compensation point, but occasionally experience brief periods of intense PAR availability known as sunflecks (Townsend *et al*., 2018). These sunflecks occur within seconds, rapidly changing the canopy light environment from shade to bright (Porcar-Castell & Palmroth, 2012). If frequent enough, sunflecks become an important source of energy, significantly boosting canopy photosynthesis (Pearcy, 1990; Murchie & Niyogi, 2011; Burgess *et al*., 2018). High photosynthetic productivity in the lower canopy layers therefore would arise from an efficient acclimation to such dynamic low-light environments by combining low respiration rates, high-light harvesting efficiency with sufficiently high quantum yield and photosynthetic capacity with optimised photoprotection relaxation to exploit sunflecks (Kromdijk *et al*., 2016; Hikosaka, 2016). Furthermore, it has been shown that bottom layers of cereal canopies have a key role in retaining N for later remobilization during grain filling (Lemaire *et al*., 2007; Sinclair & Sheehy, 1999) which may explain the high photosynthetic capacity observed in the lower layers of a wheat canopy (Burgess *et al*., 2016; Townsend *et al*., 2018).

Despite these insights it remains unclear if canopy photosynthetic potential is fully exploited or if it is restricted by physiological limitations associated with the regulation, adjustment and acclimation of the photosynthetic machinery across canopy layers (Murchie *et al*., 2018). We argue that ‘complete canopy’ photosynthesis has still not been fully exploited and has a large untapped potential to boost crop productivity. We hypothesise that identifying specific canopy layers or combinations of canopy layers will disentangle the complex link between photosynthesis, source – sink traits, phenological demands and ultimately yield. Once these associations are identified it will help breeders with the selection of more appropriate photosynthetic traits to screen crop varieties for higher yield.

## Materials and Methods

### Field site

Eleven spring bread wheat genotypes selected from the Photosynthesis Respiration Tails (PS Tails) panel from the International Maize and Wheat Improvement Center (CIMMYT) were grown at CIMMYT’s field station Campo Experimental Norman E. Borlaug (CENEB) in Ciudad Obregón, Sonora, México (27° 23’ 46’’ N, 109° 55’ 42’’ W, 38 mamsl). The genotypes were studied in three consecutive field seasons (2017-2018, 2018-2019, 2019-2020, referred hereafter as Y1, Y2 and Y3). Eight genotypes were studied in Y1 and three genotypes were added for Y2 and Y3 making a total of eleven. The genotypes were selected for their contrasting RUE expression at the vegetative and grain filling stages, grain filling flag leaf photosynthesis rates, yield, HI and a similar phenological range. Further information about genotypes studied can be found in (Robles-Zazueta *et al*., 2021). Mean temperature of the growing season (December-April) for the three years was 17.43 °C with average rainfall of 20.27 mm and incident PAR of 8.1 MJ m^−2^.

### Experimental design

A randomised complete block design with three replicates and two raised beds per plot was used in Y1 and the same experimental design with four replicates for Y2. For Y3 due to management logistics four replicates per genotype in flat beds were sown and drip irrigation was used. In all three years plants grew under yield potential conditions. Sowing dates were December 5th 2017, December 6th 2018 and December 18th 2019 for Y1, Y2 and Y3 respectively. Emergence dates were December 12th 2017, December 12th 2018 and December 26th 2019 (Y1, Y2 and Y3 respectively). Harvest dates were May 8th 2018, April 30th 2019 and May 13th 2020 (Y1, Y2 and Y3 respectively). Seed rate was ∼7 g m^−2^ for the three years and irrigation was supplied four times after emergence during the crop cycle in approximate 25-day intervals (pre-sowing, 25, 50, 75, 100 days after emergence). Plants were grown under optimal conditions in the field with pests, weed control and fertilisation management to avoid biotic and abiotic limitations to yield. In Y1 fertilization was applied in the form of urea (200 kg N ha^−1^) 25 days after emergence (DAE). For Y2 fertilization was divided in 100 kg N ha^−1^ 25 DAE and another 100 kg N ha^−1^ 58 DAE. Finally, for Y3 100 kg N ha^−1^ were applied 30 DAE and 50 kg N ha^−1^ 50 DAE; 50 kg P ha^−1^ were applied in the three cycles when the first application of N was made. A diagram summarizing the main growth stages where the different traits were measured is shown in Figure 1.

**Figure 1.**
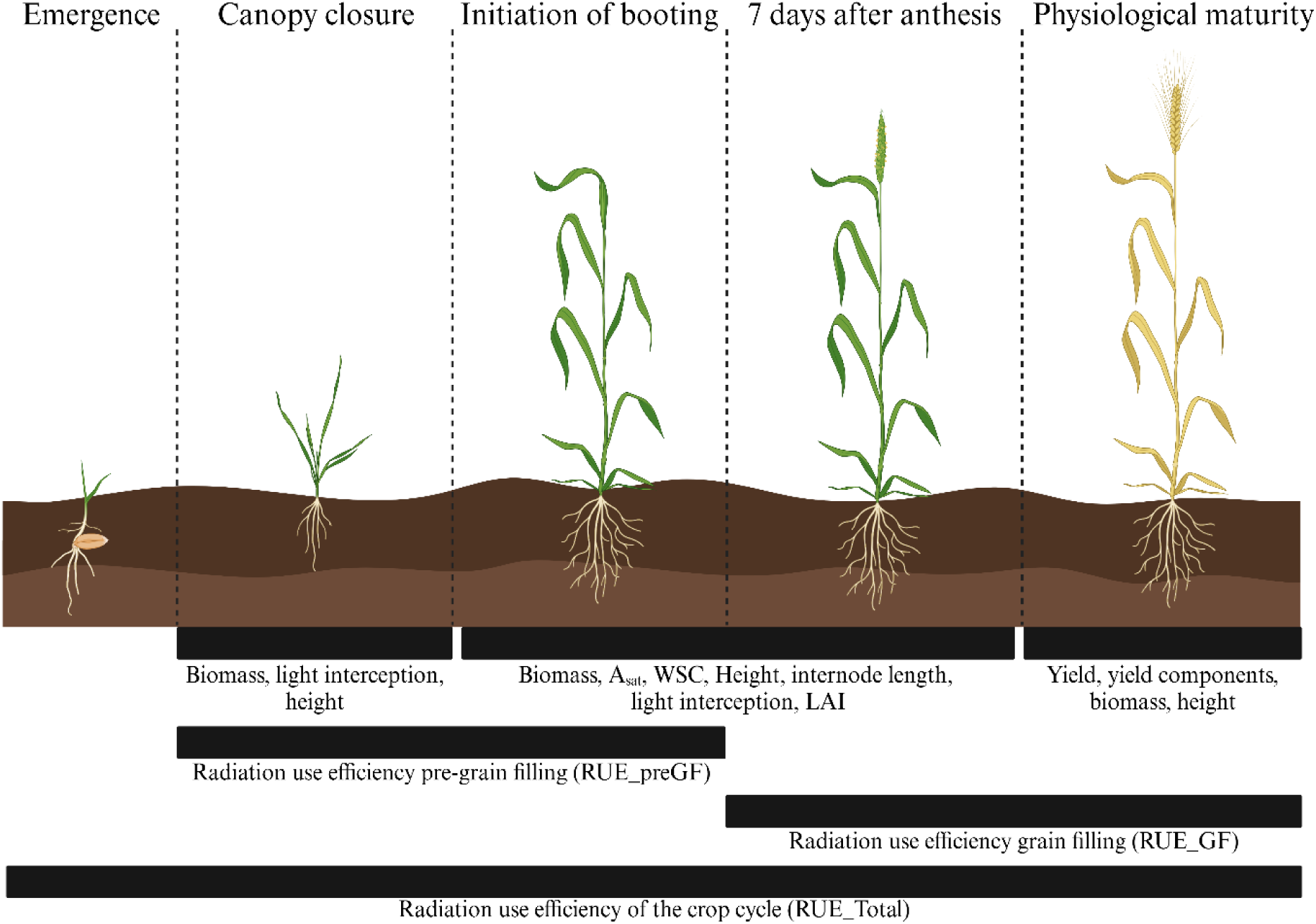
Overview of source and sink traits measured in our study. The diagram illustrates the growth stages from emergence to physiological maturity which are key for yield formation. Traits measured at each growth stage are indicated below the timeline.

### Agronomical traits

Phenological stages were scored visually according to the Zadoks growth scale for cereals (Zadoks *et al*., 1974). Growth stages recorded were initiation of booting (GS41, InB), heading (GS55, H), anthesis (GS65, A) and physiological maturity (GS87, PM).

Canopy architecture was characterized by measuring canopy height at PM in the south, north and middle areas of each bed using a measuring tape attached to a 1.5 m stick, then six values per plot were averaged for Y1 and Y2, in Y3 measurements were collected using the same protocol but only in one plot. Leaf width and length were measured at InB and A7 with a ruler in the middle area of the flag, second and third leaves. At A7 we measured leaf angles considering the ligule and curvature (point where the leaf bent) in flag, second and third leaves with a protractor and the distance from the stem to the tip of each leaf was measured using a ruler, all these data were collected from the ligule as described in (Moroyoqui-Parra *et al*., 2024). Finally, spike and awn length were measured in six shoots per plot following established field phenotyping protocols (Pask *et al*., 2013).

Light interception (LI) was measured using a ceptometer (AccuPAR LP-80, Decagon Devices, Pullman, WA, USA) at E40, InB and A7 to calculate the light intercepted and extinction coefficient by the canopy. Incident, reflected and transmitted PAR through the canopy were measured from 11:00-13:00 when clear skies and low wind speed conditions prevailed. The following equation was used to calculate the percentage of LI by the canopy:

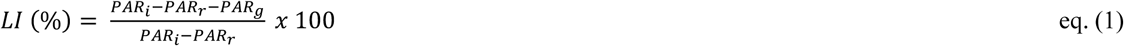

where LI (%) is the percentage of light intercepted by the canopy, PAR_i_, PAR_r_ and PAR_g_ are the incident, reflected and transmitted PAR respectively.

Twelve shoots were randomly selected for biomass partitioning where plant organs were separated from stem, green area of flag, second, third and remaining (below third leaf) leaves at InB and A7. Leaf green areas were measured using a leaf area meter (LI 3100C, Licor Biosciences, Lincoln, NE, USA). Finally, samples were dried in an oven for 2 days at 70°C, weighed and data was used to calculate the leaf area index (LAI) as follows:

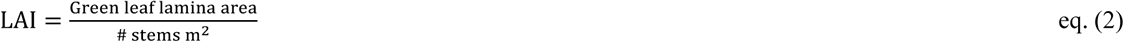

Extinction coefficient was calculated based on Beer’s law modified by Monsi and Saeki to study plant canopies (Hirose, 2005) as follows:

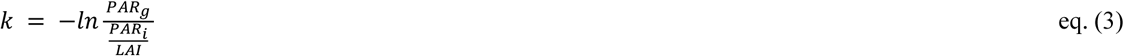

where ln is the natural logarithm, PAR_g_ the transmitted PAR, PAR_i_ the incident PAR and LAI the leaf area index of the canopy.

### Source traits

Aboveground biomass was sampled following (Robles-Zazueta *et al*., 2021). Briefly, samples of biomass at InB, A7 and PM were collected in 0.4 m^2^ (E40) and 0.8 m^2^ (InB, A7), leaving 25 and 50 cm respectively at the northern side of the plots to reduce border effects in subsequent biomass samplings. All fresh biomass was weighed, and a subsample of 50 shoots was weighted and dried in an oven at 70 °C for 48 h, to record dry weight. For biomass at PM, calculations were made from the measurement of yield components. For every growth stage, the aboveground biomass was calculated according to (Pask *et al*., 2013):

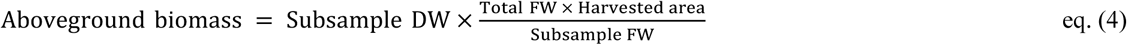

RUE was estimated from the slope of the linear regression between accumulated aboveground biomass and the corresponding accumulated intercepted PAR during the determined growth period (Monteith, 1977). Incoming radiation from a nearby meteorological station was used to calculate the accumulated PAR multiplying irradiance by 0.45 to convert it to PAR and ceptometer (AccuPAR LP-80, Decagon, Pullman, WA, USA) measurements were used to correct the accumulated radiation for the fraction of absorbed PAR of each genotype using the same approach as (Robles-Zazueta *et al*., 2021).

Photosynthesis was measured using an infrared gas analyzer (IRGA, Licor 6400 XT, Licor Biosciences, Lincoln, NE, USA) at InB (Y1 and Y2) and A7 (Y1, Y2, Y3). Spot measurements (A_sat_) were made on healthy plants using the leaf chamber fluorometer (6400-40 Licor Biosciences, Lincoln, NE, USA) in order to replicate environmental conditions from the study site (1800 µmol m^−2^ s^−1^ PAR, 28 °C for block temperature, 400 ppm CO_2_ and 60-70% relative humidity). Leaf chlorophyll content was measured using a SPAD-502 meter (Konika Minolta, Japan).

Measurements were conducted on the flag (top of the canopy), second (middle of the canopy) and third leaves (bottom of the canopy) in two main shoots per plot as indicated in (Robles-Zazueta *et al*., 2022). Measurements were performed between 10:00-15:00 as this timeframe has been found to maximize the stability and accuracy of the measurements (Evans and Santiago, 2014). Leaf photosynthesis was upscaled to canopy level by multiplying the LAI of each layer to A_sat_ values. It was found not necessary to correct for light saturation using a light response curve as in Townsend *et al*., (2018) who showed that A_sat_ was a good indicator of assimilation in all leaves (top, middle and bottom) for a PPFD range from 2000 µmol m^−2^ s^−1^ down to below 200 µmol m^−2^ s^−1^ (Supplemental Figure 1). For this study, spike and stem photosynthesis was not measured therefore was not considered in our estimations of canopy photosynthesis. Calculations were made as follows:

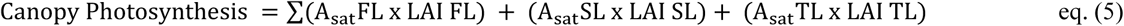

Where A_sat_ is CO_2_ assimilation under light saturated conditions (1800 µmol m^−2^ s^−1^ PAR), LAI is leaf area index, and FL, SL, TL are flag leaf, second leaf and third leaf respectively.

Water soluble carbohydrate (WSC) content was measured in stems and spikes of 12 randomly selected shoots at A7. Samples were dried in an oven at 70 °C for 48 hours and then milled for lab colorimetry analysis following the protocol described in (Pask *et al*., 2013).

### Sink traits

When the genotypes reached PM, 50 shoots were randomly harvested from each plot and dried in an oven at 70 °C for 48 hours. Then the spikes were threshed to separate the grains from the rest of biomass and the harvest index was calculated as follows:

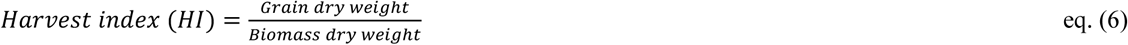

Yield was sampled in the field using an automated harvest machine (LD 350, Wintersteiger AG, Austria) and plot length was measured before yield was measured to correct for the plot area lost from the biomass sampling at previous growth stages (Pask *et al*., 2013) with a minimum area harvested of 2 m^2^ for yield estimations. From the yield sample, a subsample was collected to be processed in the lab to measure grain moisture content, to calculate the thousand grain weight (TGW), number of grains per spike (GSP), grains m^−2^ (GM2), and afterwards the grain weight per spike (GWSP) and the number of spikes m^−2^ (SM2) were calculated. Yield was calculated according to (Pask *et al*., 2013):

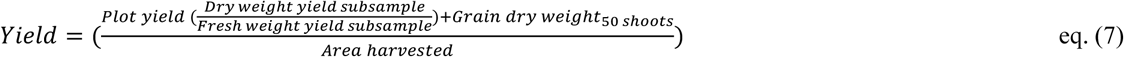

### Statistical analysis

Bilinear unbiased estimators (BLUEs) were calculated for each trait by analysing data collected in the three years using a general linear model with the R package lme4 (Bates *et al*., 2015) implemented with the graphic user interface META-R v6.04 (Alvarado *et al*., 2020). Days to InB and A, were used as covariates to correct the physiological traits for sampling variation at the vegetative and grain filling periods. To calculate BLUEs the following equation was used:

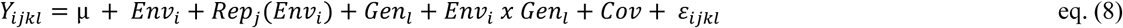

Where Y*_ijkl_* is the trait of interest, µ is the mean effect, Env*_i_* is the effect of the *i*th environment, Rep*_j_* is the effect of the *j*th replicate within the *i*th environment, Gen*_l_* is the effect of the *l*th genotype, Env*_i_* x Gen*_l_* are the effects of the *i*th environment and *l*th genotype interaction, Cov is the effect of the covariate and ε*_ijkl_* is the error associated with the environment *i*, replication *j*, *k*th incomplete block and *l*th genotype.

Broad sense heritability (H^2^) across the three years was calculated as follows:

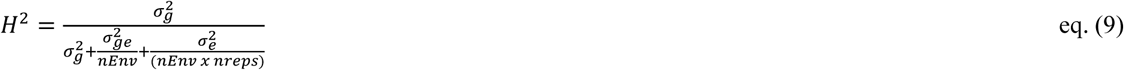

Where 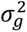 is the genotype error variance, 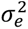 is the environment error variance, 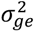 is the genotype x environment interaction error variance, nEnv is the number of environments and nreps the number of replicates.

## Results

### Leaf and canopy photosynthesis

Canopy photosynthetic rates (upscaled by multiplying each layer LAI) ranged from 84.57 µmol m^−2^ s^−1^ to 130.97 µmol m^−2^ s^−1^ at InB and 75.55 µmol m^−2^ s^−1^ to 116.94 µmol m^−2^ s^−1^ at A7 and no statistically significant differences were found (Supplemental Table 1). A similar observation was made for A_sat_ from individual layers at InB with no significant differences found between genotypes within the canopy layers (Table 2). In our study, both canopy and layer photosynthesis rates were higher at InB and decreased at A7. At InB the highest A_sat_ rates were found in the middle layer (26.91 µmol m^−2^ s^−1^) and top layer rates were statistically similar (25.58 µmol m^−2^ s^−1^) with the lowest A_sat_ rates found at the bottom layer (15.95 µmol m^−2^ s^−1^) (Figure 2A). In contrast to InB, at A7 statistically significant differences among genotypes were found in the middle and bottom canopy layers (Table 2), with a decreasing trend in A_sat_ rates from top (21.99 µmol m^−2^ s^−1^) to bottom layers (10.19 µmol m^−2^ s^−1^) of the canopy (Figure 2B).

**Figure 2.**
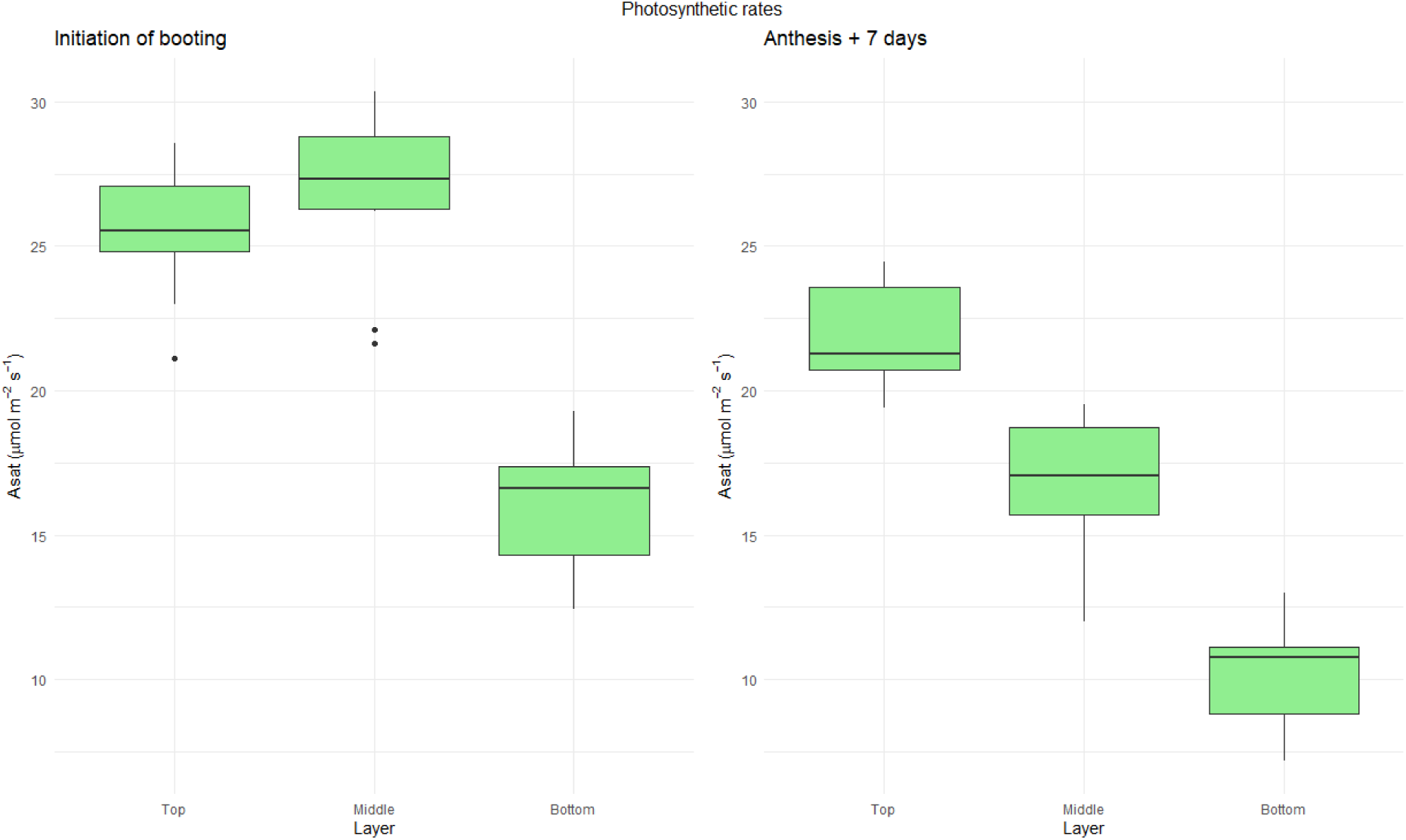
Photosynthetic rates measured at saturating light conditions (Asat) in the top, middle and bottom layers of the canopy at initiation of booting (A) and 7 days after anthesis (B).

**Table 2.**
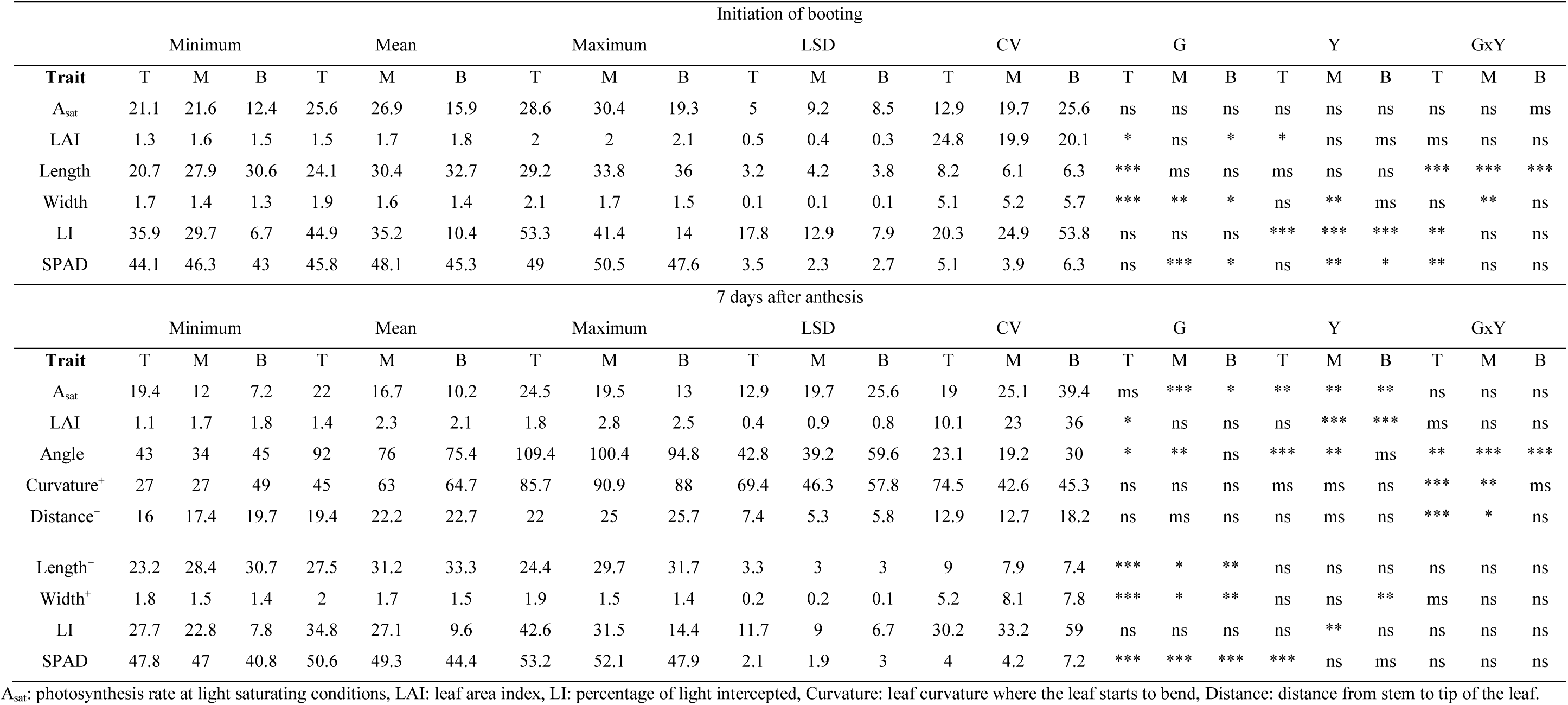
Physiological traits measured in the field study in the different canopy layers. Data presented is the mean from the field experiments from the three years, minimum, maximum, least significant differences (LSD), coefficient of variation (CV) and the statistical differences caused by genotypes (G), years of data collection (Y) or the interaction between GxY. * = significant at p<0.05, ** = significant at p<0.01, *** = significant at p<0.001, ms = significant at p<0.1, ns = no statistical significance, T = Top layer of the canopy, M = Middle layer of the canopy, B = Bottom layer of the canopy. ^+^Data collected in two years of study.

At the top of the canopy, photosynthesis is associated with architecture traits such as internode 2 and 3 length and flag leaf angle. As leaves with larger angles tend to have larger areas compared to erect leaves, there is a potential trade-off indicating that light capture may be important in the top layer, assuming that an erect leaf is likely to intercept more light over the day than a less erect one. (Figure 3A). In the middle layer photosynthesis is linked to N by percentage (N%) and by mass (SLN), suggesting that N is an important driver of photosynthesis in the middle layer where it sets up a balance between N remobilization to the spikes (before the flag leaf) and N utilization for photosynthesis (Figure 3B). Finally, angles and width are key at the bottom layer where photosynthesis is limited by efficiency of harvesting the reduced amount of light received and similarly to the middle layer SLN also plays a key role in assimilation (Figure 3C). Overall, our results show that there is a tendency across canopy positions for architectural traits to group together and layer differences in A_sat_ and N-related traits (Figure 3).

**Figure 3.**
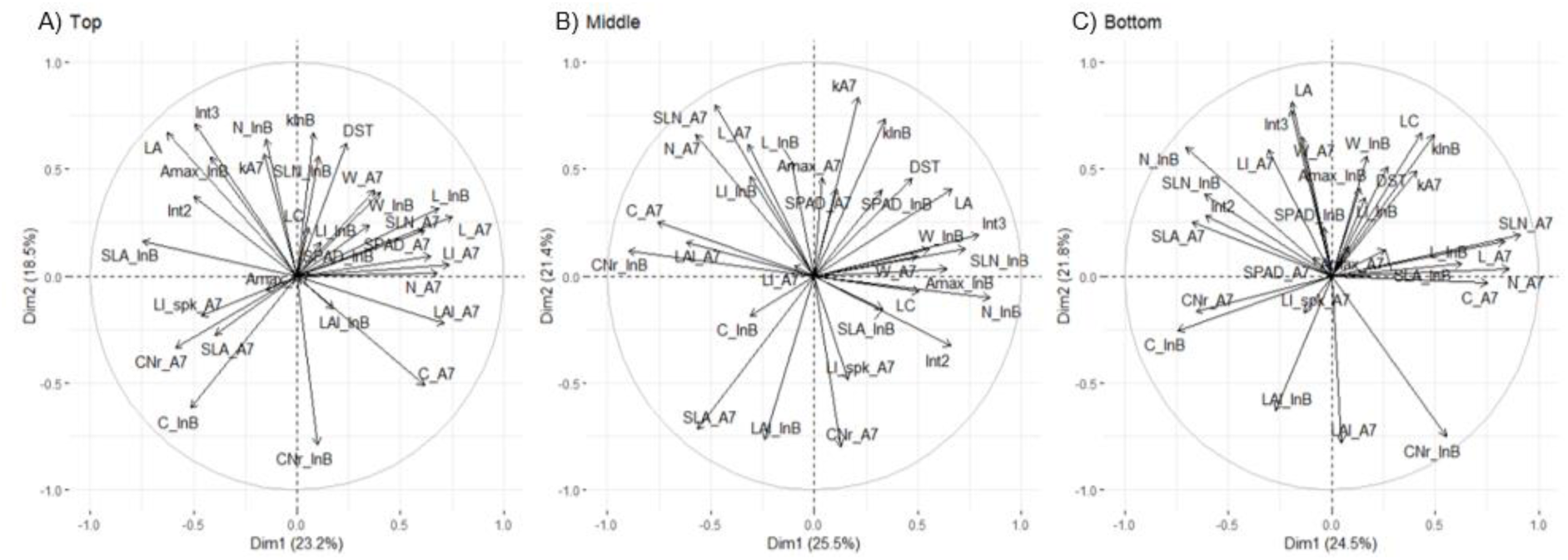
PCA biplots of photosynthesis association with traits measured in the different layers of the canopy at initiation of booting (InB) and 7 days after anthesis (A7).

### Source and sink traits

Grain yield across years was 622.6 ± 30.8 g m^−2^ (Supplemental Table 1). Yield component traits (HI, TGW, GSP, GM2 and SM2) showed highly statistically significant differences among genotypes. RUE during the vegetative period (RUE_preGF) ranged from 2.11-2.56 g MJ^−1^ and was higher compared to RUE from grain filling (RUE_GF, 0.85-1.55 g MJ^−1^) or RUE from the whole crop cycle (RUE_Total, 1.51-1.8 g MJ^−1^) (Figure 4). Differences among genotypes were identified for RUE_GF and RUE_Total at p<0.1 and p<0.05, respectively (Supplemental Table 1). Aboveground biomass ranged from 201.64 ± 33.92 g m^−2^ in E40 to 1322.83 ± 90.55 g m^− 2^ in PM (Figure 4). At PM the highest variability was found between genotypes and across years (p<0.01), but in general the sink traits varied more between genotypes and years than the source traits (Supplemental Table 1).

**Figure 4.**
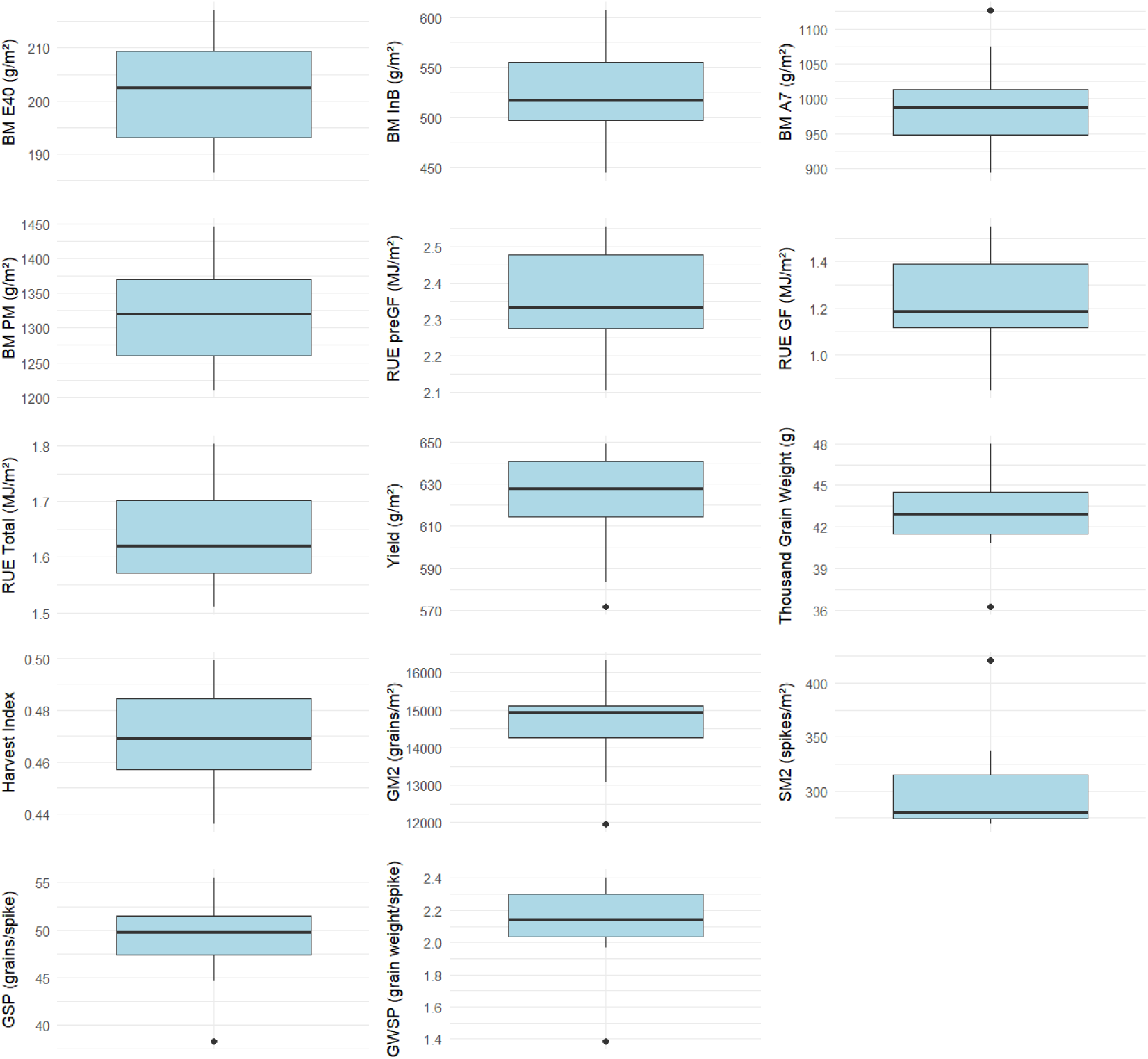
Boxplots of the main source and sink traits measured during the growth cycle. E40: 40 days after emergence, InB: initiation of booting, A7: 7 days after anthesis, PM: physiological maturity, preGF: pre grain-filling, GF: grain-filling, Total: whole crop cycle.

### Canopy photosynthesis association with source and sink traits

The strongest associations between yield and photosynthetic capacity was found in the top layer 7 days after anthesis, however the bottom layer at initiation of booting shows potential as a positive marginally significant (p<0.1) association was found, additionally, bottom of the canopy A_sat_ at InB is positively linked to water soluble carbohydrates in the spikes, HI and GSP (Supplemental Figure 3). Furthermore, canopy A_sat_ at InB is significantly associated with BM_A7, middle layer A_sat_ at InB and closely associated to BM_InB and RUE_preGF (Supplemental Figure 3).

Overall, the association between RUE measured in the different growth periods and A_sat_ measured at different layers of the canopy was weak both for A_sat_ measured during InB and A7 (Figure 5). A_sat_ measured in the top of the canopy at InB correlated significantly with RUE_GF (R^2^ = 0.43, p<0.05) (Figure 5A). When A_sat_ was measured at A7 the correlations with RUE improved (RUE_preGF and top A_sat_, p = 0.15; RUE_GF and middle A_sat_, p<0.1) and our results indicate the importance of considering different timepoints to evaluate complex associations between traits (Figure 5B).

**Figure 5.**
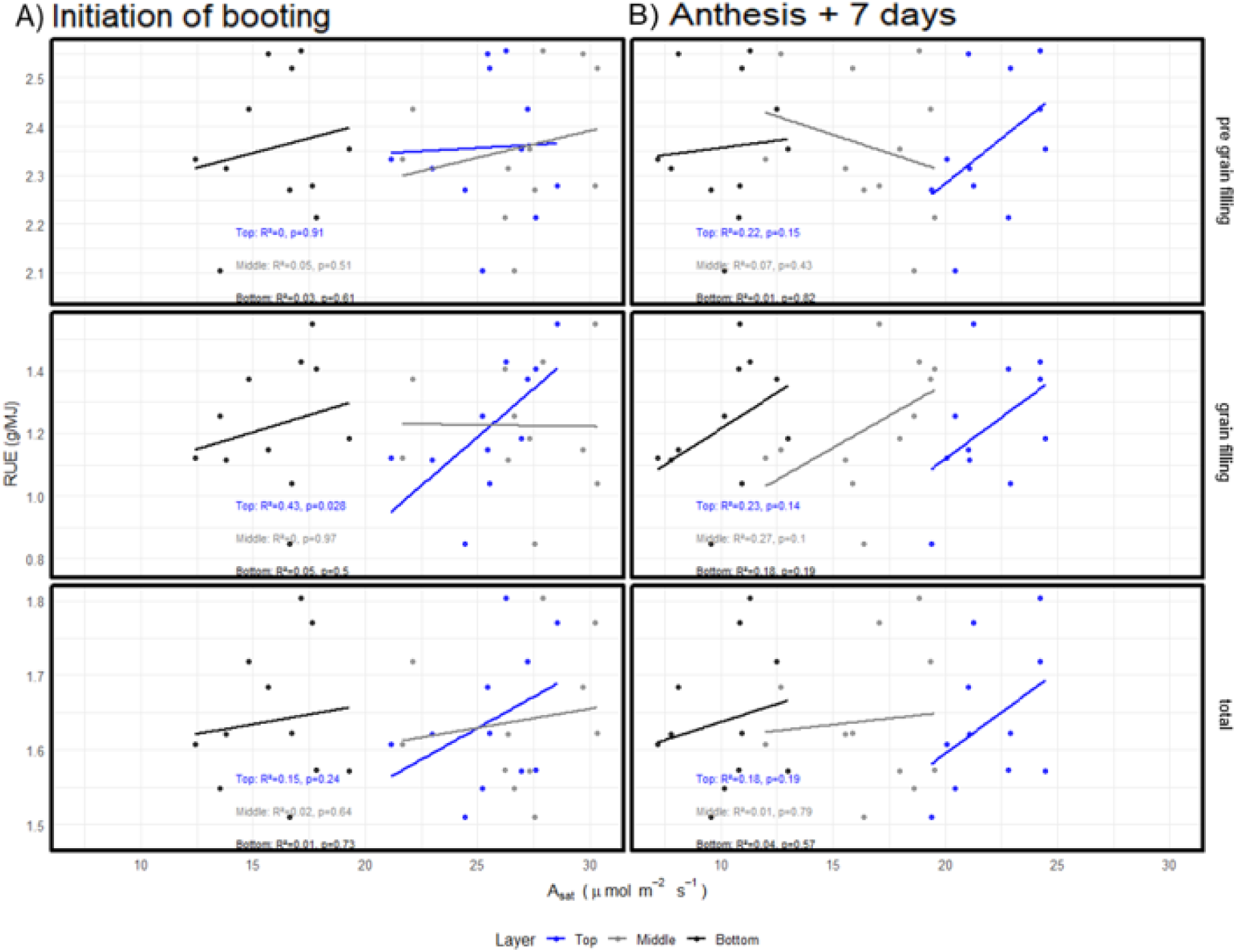
Linear regressions between radiation use efficiency and CO2 assimilation at light saturating conditions measured at different layers of the canopy at initiation of booting (A) and 7 days after anthesis (B). Blue dots: Top of the canopy, grey dots: middle of the canopy, black dots: bottom of the canopy.

When combining different layers of the canopy the relationship between RUE and A_sat_ significantly improved compared to using individual layers. RUE_preGF correlated significantly with the top + bottom layer combination (R^2^ = 0.42, p<0.05) and marginally significant correlations when the three canopy layers were considered (R^2^ = 0.26, p = 0.11) (Figure 6A). Different A_sat_ layer combinations measured at A7 correlated better with RUE. Top + bottom layers shown a significant correlation with RUE_preGF (R^2^ = 0.42, p<0.05), middle + bottom layers were positively correlated with RUE_GF (R^2^ = 0.31, p<0.1) and top + bottom layers had a marginally significant correlation with RUE_Total (R^2^ = 0.28, p<0.1) (Figure 6B).

**Figure 6.**
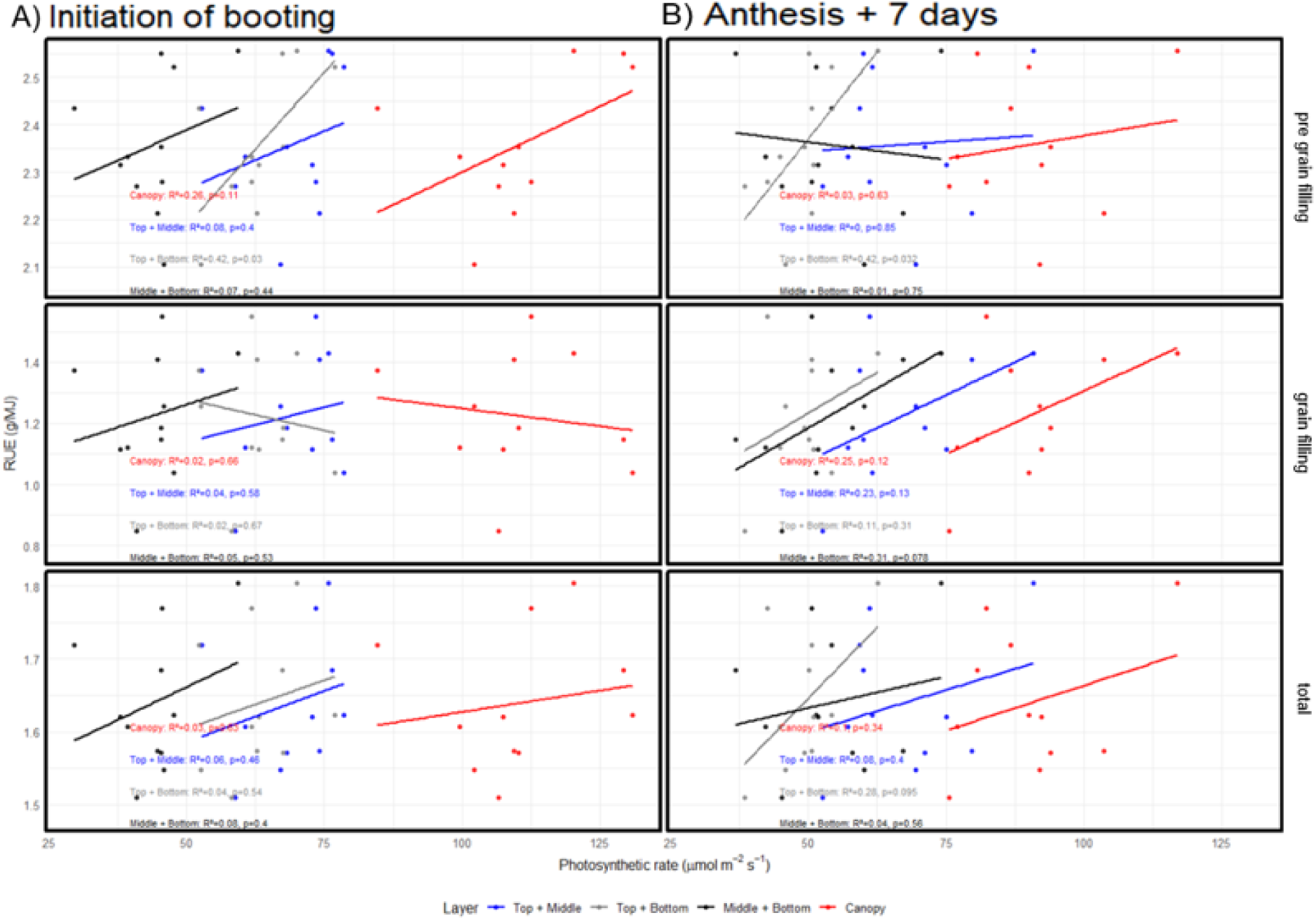
Linear regressions between radiation use efficiency and CO2 assimilation at light saturating conditions measured by combining different layers of the canopy in initiation of booting (A) and 7 days after anthesis (B). Blue dots: Top and middle, grey dots: top and bottom, black dots: middle and bottom, red dots: top, middle and bottom.

### Photosynthesis and its association with yield

First, the influence of canopy architecture on yield formation, via light interception, biomass accumulation and RUE was explored. Differences in canopy, stem and spike architectures were found across all growth stages (Supplemental Table 1). LAI showed genetic variability at canopy (Supplemental Table 1) and individual leaf layers at InB (Table 2) with no differences found at A7. Similarly, statistically significant differences were found in leaf angles measured in the top and middle layers of the canopy (Table 2). Despite the differences in canopy architecture, all the genotypes captured the same amount of light in all the growth stages studied evidenced by the lack of significance in the LI GxYxGS interaction (Supplemental Table 2).

Biomass accumulation throughout the growing season is a key component for yield formation, and its final expression at physiological maturity is an important breeding target. Our results showed that A_sat_ at InB from the top layer is associated with increased biomass at canopy closure (R^2^ = 0.28, p<0.1) and the middle layer assimilation at A7 is associated with BM_A7 (R^2^ = 0.32, p<0.1) (Supplemental Figure 2).

Significant relationships were found between photosynthetic rates and yield at InB and A7. A_sat_ at InB from the bottom layer of the canopy had a positive association with yield (R^2^ = 0.33, p<0.1) and the top layer of the canopy shown a positive association as well (R^2^ = 0.24) albeit not significant (Figure 7A). At 7 days after anthesis, a strong positive link between yield and A_sat_ was found (R^2^ = 0.54, p<0.01), and the bottom layer also showed a positive trend however not significant (R^2^ = 0.23) (Figure 7B).

**Figure 7.**
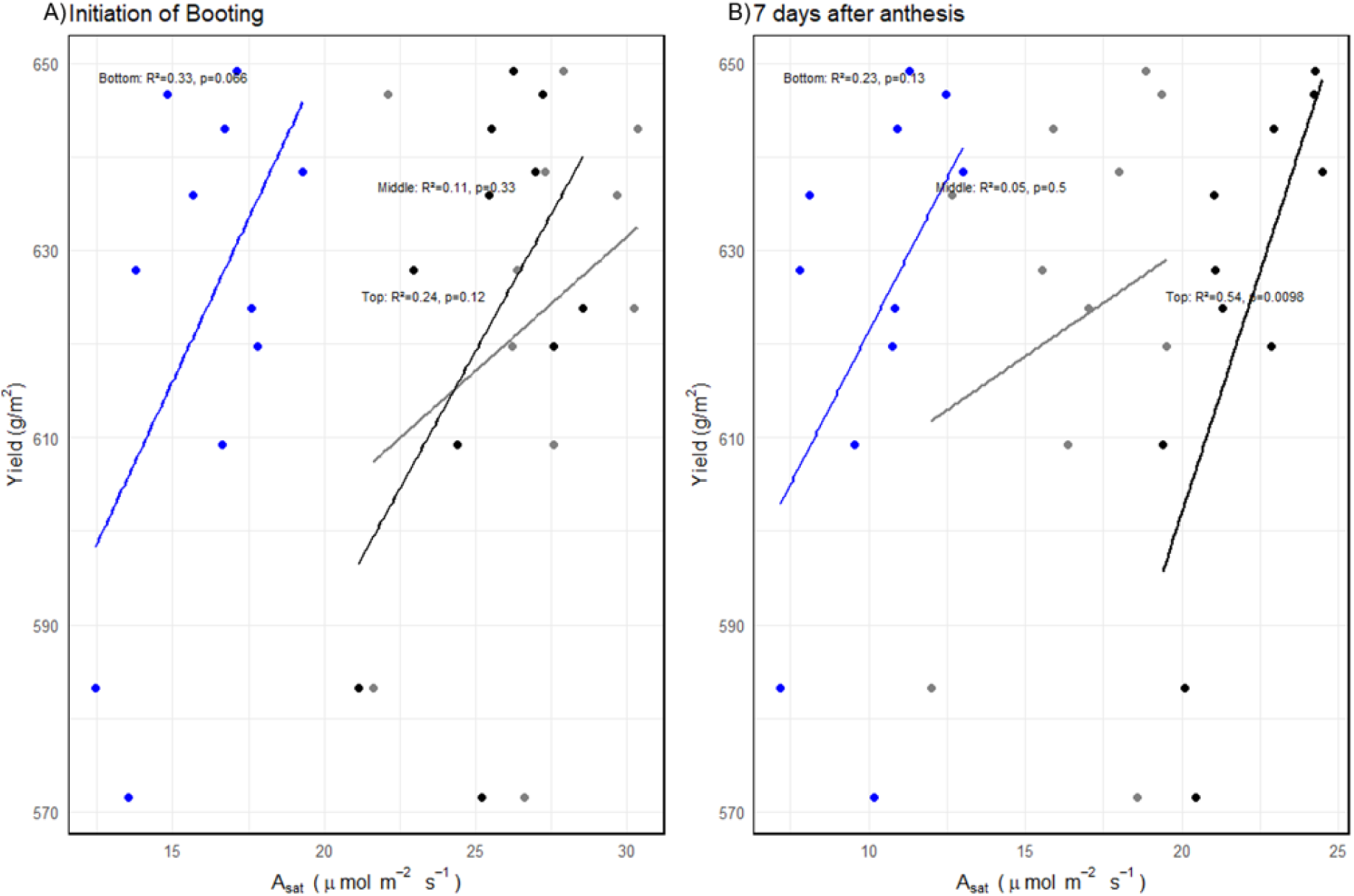
Linear regressions between grain yield and CO2 assimilation at light saturating conditions at initiation of booting (A) and 7 days after anthesis (B). Black dots: top, grey dots: middle, blue dots: bottom.

Combining different canopy layers revealed that i) they did not generally improve the above relationships compared to the ones established with individual layers and ii) when all layers were combined into total canopy photosynthesis, the relationship with yield became weaker still either at InB nor A7 (Figure 3). Top + bottom layers photosynthesis in both InB and A7 were found to associate with yield significantly (R^2^ = 0.28, p<0.1 Figure 8A; R^2^ = 0.38, p<0.05 Figure 8B, respectively).

**Figure 8.**
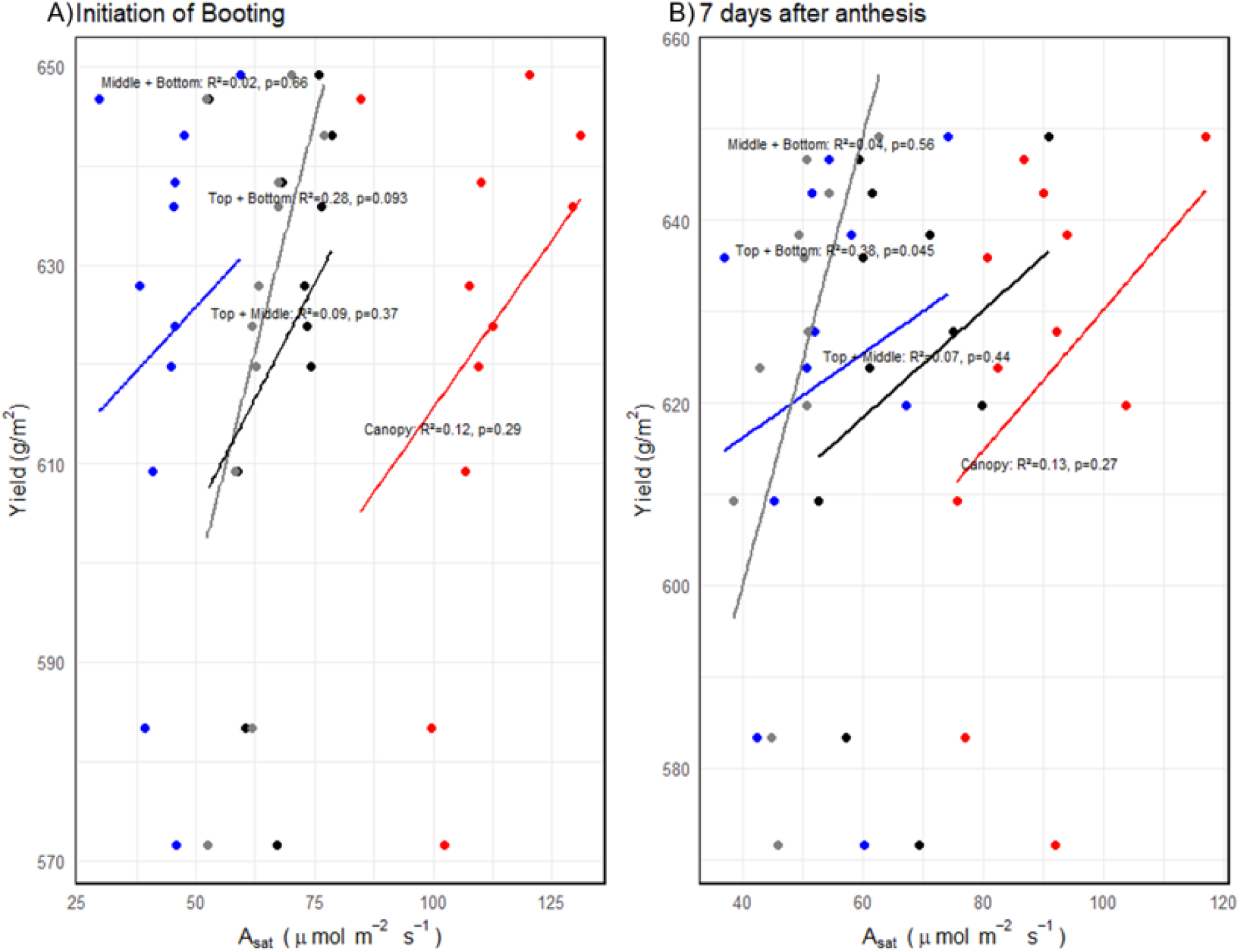
Linear regressions between grain yield and CO2 assimilation at light saturating conditions measured by combining different layers of the canopy in initiation of booting (A) and 7 days after anthesis (B). Black dots: top and middle layer, grey dots: top and bottom layer, blue dots: middle and bottom layer, red dots: combined layers.

## Discussion

### The overlooked role of shaded leaves in crop yield formation

This is the first study to consider the impact of photosynthesis at discrete layers of wheat canopies on biomass, RUE and yield in wheat; as well as the combined impact of different layer combinations. Our results suggest that the relationship between yield, biomass and RUE with leaf photosynthesis has not been consistent due to the lack of consideration of the contribution of photosynthesis within different canopy layers. Net photosynthesis is likely to be affected by differences in the canopy micro-environment and by developmental progression through phenological stages.

We confirm that understory photosynthesis plays an important role in yield formation, despite the fact that we only considered photosynthesis under saturating light, where it would apply to sunflecks during which photosynthetic capacity could be reached briefly after a period of induction. If there was sufficient sunlight penetration through the canopy this would increase the value of maintaining high A_sat_ in lower layers. Recent evidence using wheat canopies with varying canopy architecture showed that upright canopies with better light distribution had greater yield (Richards *et al*., 2019; Song *et al*., 2017b). Light response curves of wheat leaves at different canopy layers showed that A_sat_ was a good indicator of assimilation in all leaves (top, middle and bottom) for a PPFD range from 2000 µmol m^−2^ s^−1^ down to below 200 µmol m^−2^ s^−1^ (Townsend *et al*., 2018). Therefore, from our results we can conclude that A_sat_ in different canopy layers may be sufficient to select wheat varieties with higher yield as this trait is a good indicator of canopy photosynthesis.

What are the optimal characteristics of lower canopy leaves? Our work contrasts with the previous study from (Townsend *et al*., 2018) who studied canopies with high LAI and concluded that sunflecks were relatively rare and brief at the bottom of dense wheat canopies and suggested that even though A_sat_ rates were high due to high leaf N availability, induction rates would be too slow to fully exploit high light availability. That being the case, the importance of using A_sat_ for phenotyping would be diminished in dense wheat canopies. In lower canopy layers with low light availability it is crucial to consider specific leaf characteristics such as fast induction of photosynthesis (Taylor & Long, 2017), light harvesting efficiency and mitochondrial respiration (Zhu *et al*., 2018; Scafaro *et al*., 2021). Our data shows that during grain filling, associations between A_sat_ in middle and lower layers with grain yield are low, presumably due to the high photosynthesis levels in middle and bottom layers before anthesis to determine higher grain number and during grain filling top layer photosynthesis is higher to maintain feeding the grain as photosynthesis in modern cultivars is sink limited. It may be advantageous therefore to retain high A_sat_ for as long as possible if this does not interfere with N remobilization.

### Addressing the lack of consensus in photosynthesis and yield improvement

Historically, the relationship between gas exchange traits (including light saturated photosynthetic CO_2_ assimilation [A_sat_], stomatal conductance [*gs*], dark respiration [R_d_]) and yield have been inconsistent (Supplemental Table 3). This may be because single steady state measurements do not capture other key yield – forming criteria such as dynamic photosynthesis, canopy gradients and sink limitation. Carmo-Silva and colleagues (2017) found positive statistically significant correlations with HI and yield when flag leaf assimilation was measured at 1000 µmol m^−2^ s^−1^, which in tropical or subtropical latitudes are probably at the upper end of radiation levels expected to be measured in the middle layer of wheat canopies. Interestingly flag leaf A_sat_ (1800 µmol m^−2^ s^−1^) did not correlate with HI or yield perhaps because these conditions did not adequately match field values (Driever *et al*., 2014). Nonetheless consistent relationships in most studies which focus only in A_sat_ at the top layer are not found. The most common approach has been modelling or flux chambers (Zhu *et al*., 2012; Song *et al*., 2016; Wu *et al*., 2018) but these do not consider each individual leaf layer within the canopy and their empirical contribution to important agronomic traits such as biomass accumulation or yield.

Another key factor in the relationship between photosynthesis and yield is phenology: leaf photosynthesis (as measured by brief snapshots) is expected to have a greater contribution at key phases of growth (Murchie et al., 2023; Reynolds et al., 2022). It is notable that phenology influenced the strength of the relationships between performance traits and photosynthesis in the current study, with the lower layers being more important during booting, prior to N remobilisation. This is consistent with this phase being important for the synthesis of stem carbohydrate. The onset of senescence and N remobilisation during grain filling renders the role of photosynthesis in lower layers less critical.

Previous studies that have explored the correlations of source and sink traits with photosynthesis have focused on flag leaves (wheat, rice) or top of the canopy leaves (cassava and sorghum) and there is a range of environments (yield potential, drought, different N fertilisation rates, glasshouse and growth chamber) that have an effect on the source-sink ratios of plants, which adds a confounding effect to the study of these relationships. In our study, A_sat_ expressed in the middle layer of the canopy, both at InB and A7, were best associated with genotypic differences in final biomass (r^2^ = 0.18, 0.25, respectively) as flag leaves are for most of the crop cycle subjected to light saturating conditions. The relationship with biomass in this study is within the range of previous studies for sorghum at panicle initiation (r^2^ = 0.32) (Peng *et al*., 1991), wheat before anthesis (r^2^ = 0.25) (Gent, 1995), but smaller than the relationships presented for wheat grown in the field under drought conditions (Wada *et al*., 1994), heat stressed (Gutiérrez-Rodrı’guez *et al*., 2000; Reynolds *et al*., 2000), contrasting N fertilization regimes (Huang *et al*., 2016) and different N treatments at pre and post-anthesis (Gaju *et al*., 2016) .

On the other hand, yield and A_sat_ relationship is more established than the biomass and A_sat_ relationship. In our study photosynthetic rates at InB in every layer measured had significant correlations with yield (Supplemental table 1). Under drought and yield potential conditions, field grown sorghum showed correlations at different growth stages with yield, especially in the mid-development stage (r^2^ = 0.86, p<0.01; r^2^ = 0.83, p<0.01, for yield potential and drought, respectively) (Peng *et al*., 1991), and for wheat in the vegetative stage differences between the relationships under yield potential and drought conditions were strongly contrasting (r^2^ = 0, r^2^ = 0.4, p<0.01, respectively) (Wada *et al*., 1994).

For most of field trials that previously reported relationships between yield and photosynthesis, the strongest relationships were found at grain filling (Gent, 1995; Fischer *et al*., 1998; Reynolds *et al*., 2000; Zheng *et al*., 2011; Gaju *et al*., 2016), similarly to our results. This evidence suggests that yield improvement in wheat might has come hand to hand with increments in A_sat_ and *gs*, because selection of new varieties with greater biomass is thought to inadvertently come with greater gas exchange rates, this indicates that it will be easier to find wheat varieties with higher yield if photosynthetic rates are high during grain filling.

In contrast to field trials, when plants were studied in controlled settings the strongest relationships were found. Examples of this is rice grown under elevated [CO_2_] (Sakai *et al*., 2006), wheat grown with drought and high N fertilisation rates (Barbour and Kaiser, 2016). Spike photosynthesis of wheat grown in glasshouse with optimum irrigation showed contrastingly better relationship with yield (Zhou *et al*., 2016; Elazab *et al*., 2021) compared to spike photosynthesis measured in the field under yield potential (Molero & Reynolds, 2020); and a similar trend was found for flag leaf A_sat_ and A_max_ when wheat was analysed in growth chamber conditions (Driever *et al*., 2017) compared when the same genotypes were studied in the field (Driever *et al*., 2014) (Supplemental Table 3).

From the abovementioned examples it is clear that the range of growing conditions plants are subjected have a clear effect on the relationship between yield and photosynthesis. Since photosynthetic traits can be environmental, time specific and developmental stage dependant (Flood *et al*., 2016), this needs to be considered when screening lines in different environments if yield is to be boosted. Plants developing in controlled environments typically have higher growth rates, higher leaf N concentration (which is key for photosynthesis) and smaller leaf area which affect the plant source-sink balance (Poorter *et al*., 2016) and could explain the higher relationships found in controlled settings compared to the field (Supplemental Table 3).

### Canopy photosynthesis as a driver of yield improvement

Our results indicate that preferential improvement in the middle and bottom layers of canopy photosynthesis are most likely to boost yield. However, given that these results were derived under light saturated conditions at all canopy layers, modifications to canopy architecture are likely needed to improve light penetration through middle and bottom layers (e.g. erect leaves, increasing lower internode length, smaller LAI at the top and larger LAI at the bottom the canopy) which can potentially translate to higher plants, but recent evidence suggests that is not necessarily the case (Rivera-Amado *et al*., 2020).

It has been suggested that exploiting the genetic variation in biomass (Aisawi *et al*., 2015) and RUE (Joynson *et al*., 2021) can be an important avenue for yield improvement, and furthermore help to understand the genetic basis of physiological traits related to yield (Molero *et al*., 2019). Nevertheless, if physiological variables do not show a relationship with growth analysis, RUE or yield new physiological traits will not be introduced in breeding pipelines, luckily our study shows evidence of top, middle and bottom layer photosynthesis relating with yield.

In this study we establish that adding middle and bottom layers of the canopy to physiological studies, can help us to find genotypes with higher RUE rates, biomass and yield, as relationships of photosynthesis with RUE, biomass and yield, and *gs* with biomass were found (Supplemental table 1). Recent research has found mixed results on whether yield is source or sink limited, with indications that increments in grain filling source capacity including higher LAI and spike or leaf photosynthesis will improve yield (Rivera-Amado *et al*., 2020). On the other hand, (Quintero *et al*., 2018) found that increasing the sink size will boost yield. These mixed results suggest a source-sink co-limitation of yield; therefore, it will be paramount to increase canopy photosynthesis with an emphasis on middle and bottom layer at InB to allow plants to have greater photoassimilates reserves stored in the stems when remobilization starts at grain filling and leaf senescence at the bottom of the canopy diminishes the overall photosynthetic rates.

Exploiting canopy photosynthesis for crop adaptation to specific Target Population Environments (TPEs) represents the next frontier for yield improvement across diverse wheat megaenvironments (MEs). Wheat MEs differ considerably in temperature, radiation, water, and nutrient availability, each influencing canopy-level physiological responses. For example, optimizing canopy photosynthesis in high-radiation, irrigated MEs can be achieved by focusing on improving canopy architecture traits that maximize lower canopy illumination, such as erect leaf angles or reduced upper canopy LAI. In contrast, for moisture-limited MEs, productivity gains can be achieved by enhancing photosynthetic performance under intermittent and sub-saturating light conditions through rapid induction response, improved light harvesting efficiency in PSII, or reduced mitochondrial respiration in lower canopy layers. Additionally, in MEs experiencing heat stress during grain filling, enhancing heat and light tolerance at the canopy’s upper layers by improving non-photochemical quenching (NPQ), coupled with maintaining high photosynthetic rates in middle and lower canopy layers, may bolster stress resilience and optimize source-sink balance through effective nitrogen remobilization. Thus, targeted selection and phenotyping of canopy photosynthetic traits tailored specifically to environmental stressors and resource availability within each MEs offer breeders a robust strategy to leverage genetic diversity and achieve sustainable yield improvements.

## Conclusions

To our knowledge, this is the first effort to add different layers of the canopy to the study of the relationships between source and sink traits with gas exchange in staple crops. Our results indicate that the growth stage where measurements take place are crucial to characterize the link between photosynthesis and yield. These measurements coupled with high-throughput phenotyping methods, for example based on optical remote sensing (Furbank *et al*., 2021; Robles-Zazueta *et al*., 2022) will increase the feasibility of including photosynthetic traits into breeding pipelines for large trial screenings.

Discrepancies in the literature related to the link between photosynthetic traits and yield or biomass appear to be related to differences in growing conditions that still obscure these relationships. Future studies should consider measuring different wheat genotypes in multi-environmental trials coupled with high-throughput phenotyping of different canopy layers with modelling approaches considering the addition of spike and stem photosynthesis in at least one vegetative and one reproductive stage to catch the variability caused by phenology, environmental and management conditions where wheat is grown.

## Acknowledgements

CARZ acknowledges funding for his PhD studies (CVU 626989) from the Secretaría de Ciencia, Humanidades, Tecnología e Innovación (SECIHTI, Mexico). CARZ, GM and MPR acknowledge funding from the International Wheat Yield Partnership (IWYP) and from the Sustainable Modernization of Traditional Agriculture (MasAgro), an initiative from the Secretariat of Agriculture and Rural Development (SADER) from Mexico and CIMMYT. EHM, GM and MPR received funding from the Newton-Mexico BBSRC partnership award (grant BB/S012834/1). CARZ and MPR acknowledge funding from the Heat and Drought Wheat Improvement Consortium (HeDWIC) an initiative from the Foundation for Food and Agricultural Research (FFAR) while writing this manuscript. We gratefully thank the members of CIMMYT physiology group for their support with field trials, especially Margarita Guerra for her support with gas exchange measurements, Carolina Rivera, Francisco Pinera-Chavez and Jazmin Rodriguez for their help coordinating yield measurements and Julio César Rodríguez (Universidad de Sonora) for lending us an IRGA during the second-year field trial.

## Author contributions

**Carlos A. Robles-Zazueta:** Conceptualization, Methodology, Investigation, Project Supervision, Data Collection, Data Analysis, Visualization, Writing-Original Draft, Writing-Review and Editing; **Gemma Molero**: Conceptualization, Methodology, Project Supervision, Writing-Review and Editing; **Matthew P. Reynolds**: Funding Acquisition, Writing-Review and Editing; **Erik H. Murchie**: Conceptualization, Project Supervision, Funding Acquisition, Writing-Review and Editing.

## Data availability statement

All data used in this study is available as supplemental material both in its raw and analysed formats.

