## Supplementary Figures for "Yield from the shadows: beyond top layer photosynthesis to enhance crop productivity"

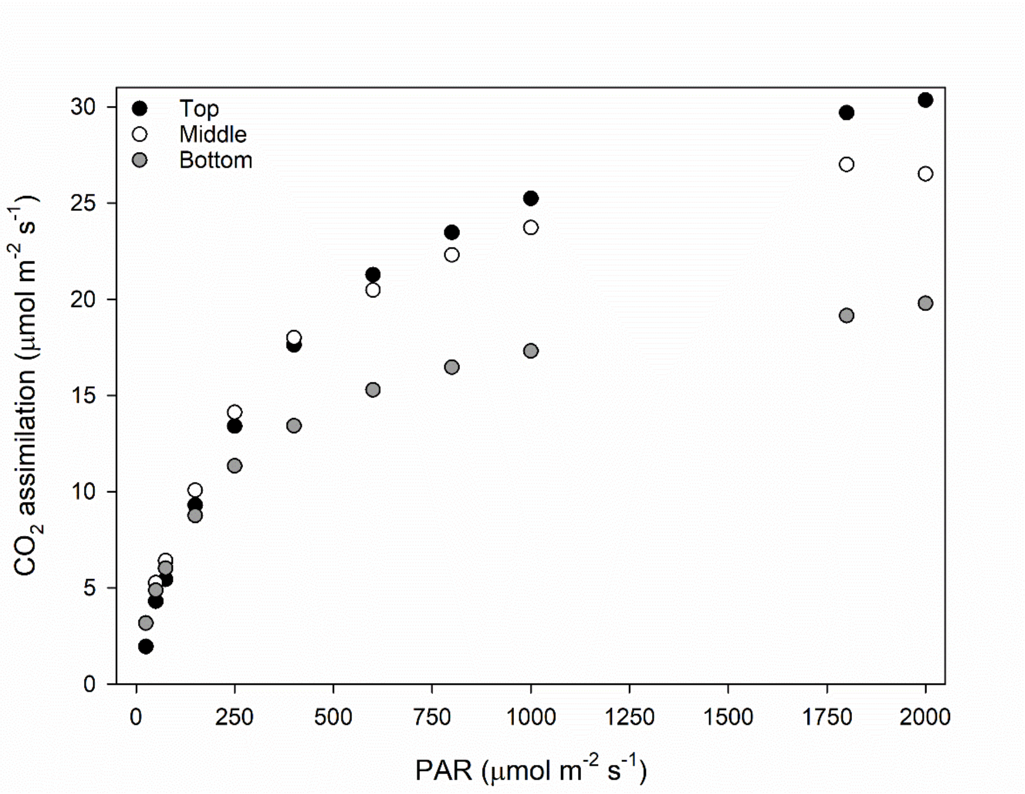


Supplemental Figure 1. Light response curves from the top, middle and bottom canopy layers. We observe that the three layers saturate around 1750 µmol m$^{-2}$ s$^{-1}$. Our $A_{sat}$ measurements were collected at 1800 µmol m$^{-2}$ s$^{-1}$ to ensure measuring maximum photosynthetic capacity at each canopy layer.

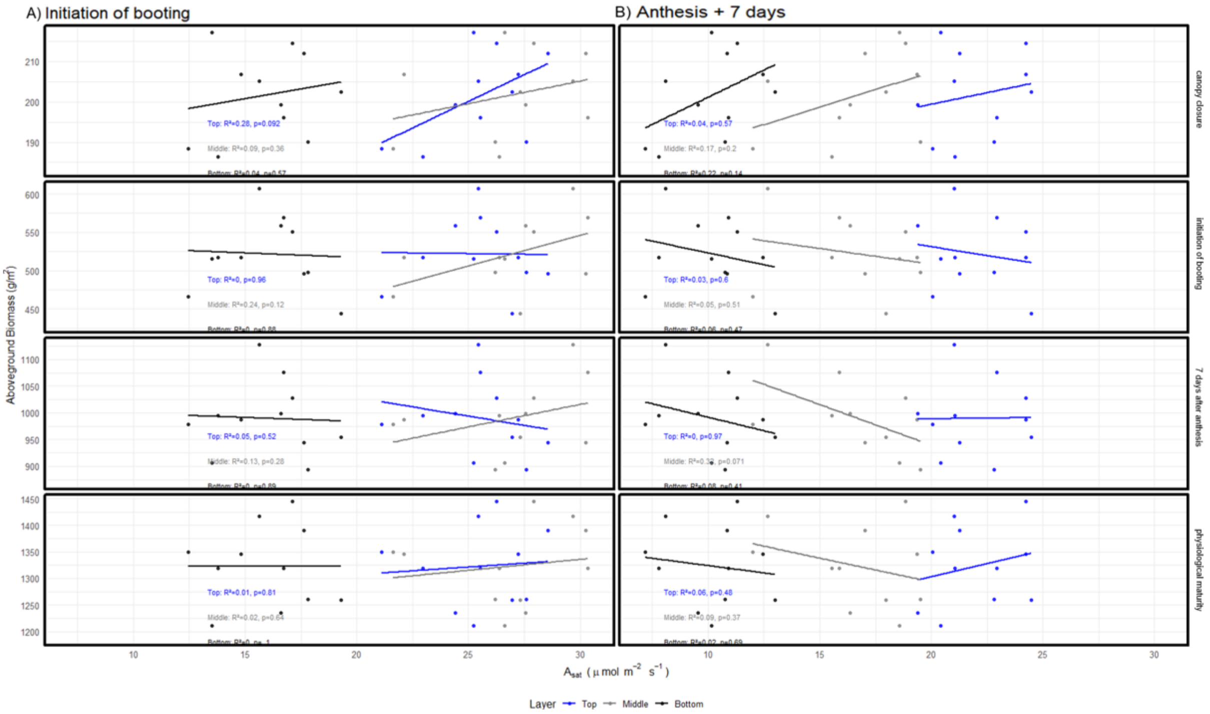


Supplemental Figure 2. Linear regressions between aboveground biomass measured 40 days after emergence, initiation of booting, 7 days after anthesis and physiological maturity with $CO_2$ assimilation at light saturation conditions measured by combining different layers of the canopy in initiation of booting (left panels) and 7 days after anthesis (right panels). Black dots: top and middle layers, white dots: top and bottom layers, grey dots: middle and bottom layers.

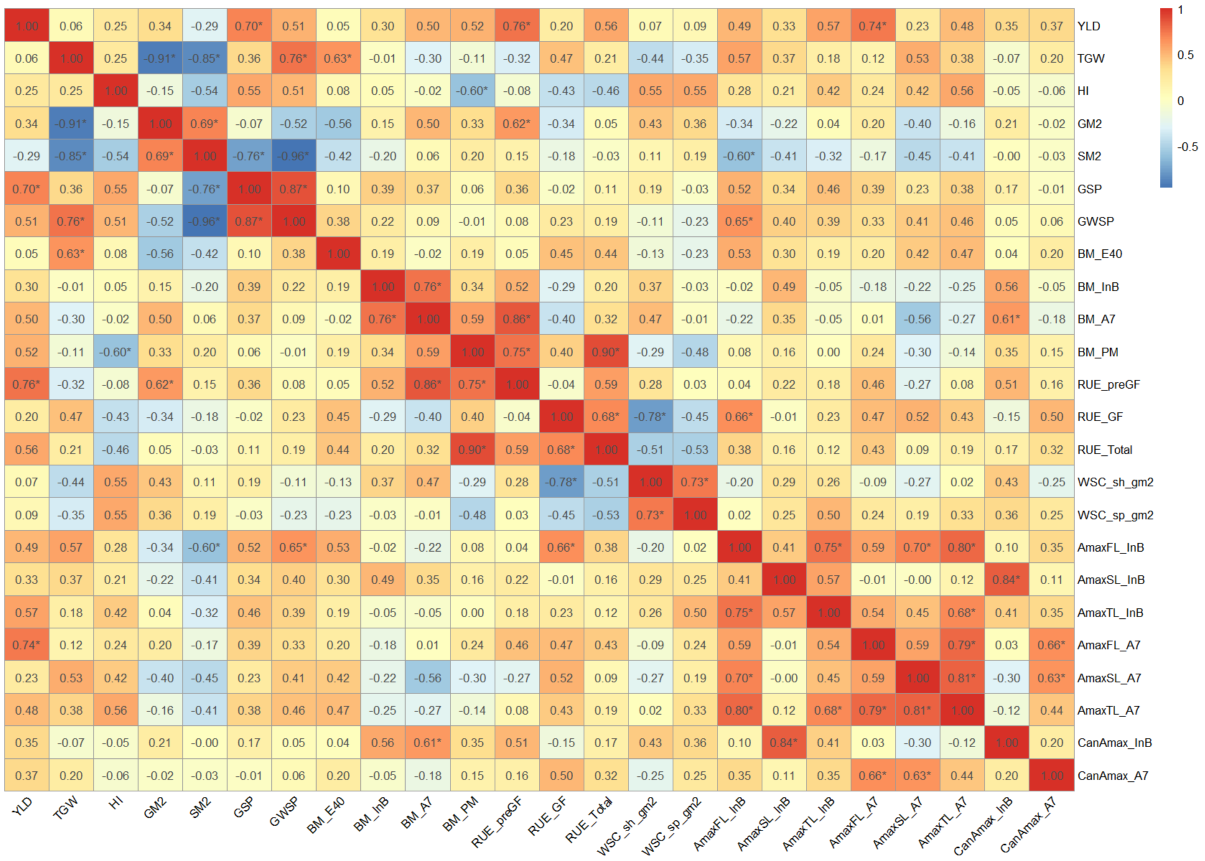


Supplemental Figure 3. Heatmap correlation matrix showing associations between source and sink traits. Colour scales ranging from blue (negative) to red (positive) correlations indicating the magnitude of the correlation. Numeric values inside the cells represent the Pearson moment correlation coefficient and asterisks indicate significant association between traits at $p<0.05$.
