## Supplementary Tables for "Yield from the shadows: beyond top layer photosynthesis to enhance crop productivity"

Supplemental Table 1. Traits measured at plot level during 3 field seasons. Data presented is the mean (standard deviation) from 8 genotypes studied in three years plus 3 more genotypes studied two years, minimum, maximum, least significant differences (LSD), coefficient of variation (CV) and the statistical differences caused by genotypes (G), years of data collection (Y) or the interaction between GxY. * = significant at p<0.05, ** = significant at p<0.01, *** = significant at p<0.001, ms = marginally significant (0.1>p>0.05), ns = no statistical significance.

| Trait | Mean (SD) | Minimum | Maximum | LSD | CV | G | Y | GxY |
|---|---|---|---|---|---|---|---|---|
| Yield (g $m^{-2}$) | 622.6 (30.82) | 571.53 | 649.12 | 51.36 | 5.31 | * | ns | ** |
| HI | 0.47 (0.018) | 0.436 | 0.499 | 0.02 | 4.96 | *** | *** | ns |
| TGW (g) | 43.11 (1.66) | 36.24 | 47.98 | 2.88 | 6.33 | *** | *** | ns |
| GSP (# $spike^{-1}$) | 49.12 (4.37) | 38.34 | 55.53 | 5.79 | 12.69 | *** | * | ns |
| GWSP (g $spike^{-1}$) | 2.11 (0.17) | 1.39 | 2.4 | 0.26 | 11 | *** | * | ns |
| GM2 (grains $m^{-2}$) | 14604.68 (972.02) | 11963.71 | 16314.09 | 1777.86 | 8.13 | *** | *** | ** |
| SM2 (spikes $m^{-2}$) | 303.41 (32.71) | 269.4 | 420.8 | 40.25 | 12 | *** | * | ns |
| BM_E40 (g $m^{-2}$) | 201.64 (33.92) | 186.36 | 217.15 | 52.57 | 16.67 | ns | ns | ** |
| BM_InB (g $m^{-2}$) | 521.84 (60) | 444.28 | 607.09 | 98.08 | 13.4 | ms | ns | * |
| BM_A7 (g $m^{-2}$) | 989.59 (126.72) | 893.31 | 1127.18 | 137.86 | 14.23 | * | ns | ns |
| BM_PM (g $m^{-2}$) | 1322.83 (90.55) | 1210.31 | 1445.18 | 131.68 | 7.12 | ** | ** | * |
| RUE_preGF (g $MJ^{-1}$) | 2.36 (0.36) | 2.111 | 2.56 | 0.39 | 17.26 | ns | ns | ns |
| RUE_GF (g $MJ^{-1}$) | 1.23 (0.5) | 0.85 | 1.55 | 0.49 | 41.01 | ms | ns | ns |
| RUE_Total (g $MJ^{-1}$) | 1.64 (0.14) | 1.51 | 1.8 | 0.19 | 8.97 | * | *** | ms |
| Canopy $A_{sat}$_InB (μmol $m^{-2}$ $s^{-1}$) | 110.26 (24.79) | 84.57 | 130.97 | 48.92 | 21.51 | ns | ns | ns |
| Canopy $A_{sat}$_A7 (μmol $m^{-2}$ $s^{-1}$) | 90.09 (30.87) | 75.55 | 116.94 | 35.3 | 35.59 | ns | ns | ns |
| WSC_stems (g $m^{-2}$) | 122.91 (34.63) | 72.31 | 168.9 | 64.05 | 34.04 | ** | ns | ** |
| WSC_spikes (g $m^{-2}$) | 39.24 (12.79) | 27.84 | 49.62 | 18.74 | 38.49 | * | ns | ns |
| Height_PM (cm) | 110.77 (2.06) | 100.91 | 116.99 | 3.35 | 1.92 | *** | *** | ** |
| Internode 2 length (cm) | 21.25 (0.81) | 19.82 | 23.39 | 1.43 | 3.63 | *** | ns | *** |
| Internode 3 length (cm) | 13.77 (0.54) | 10.54 | 15.92 | 1.08 | 4.3 | *** | *** | *** |
| Peduncle length (cm) | 38.42 (1.34) | 35.07 | 42.73 | 2.84 | 4.05 | *** | *** | *** |
| Spike length (cm) | 11.44 (0.4) | 9.3 | 12.94 | 0.74 | 3.76 | *** | ns | *** |
| Awns length (cm) | 6.27 (0.36) | 4.84 | 6.84 | 0.35 | 5.92 | *** | *** | ns |
| Shoots_E40 (# $m^{-2}$) | 805.38 (123.45) | 710.39 | 928.65 | 128.12 | 16.98 | *** | ns | ns |
| Shoots_InB (# $m^{-2}$) | 534.65 (88.27) | 429.91 | 697.84 | 91.31 | 17.77 | *** | *** | ns |
| Shoots_A7 (# $m^{-2}$) | 454.02 (62.92) | 373.66 | 604.8 | 71.47 | 16.66 | *** | ns | ns |
| LI_E40 (%) | 89.57 (3.78) | 87.14 | 91.28 | 4.6 | 4.77 | ns | * | ns |
| LI_InB (%) | 96.4 (1.84) | 92.81 | 98.62 | 3.51 | 2.45 | ms | ms | ns |
| LI_A7 (%) | 97.7 (0.9) | 96.77 | 98.51 | 1.2 | 1.15 | ms | ** | ns |
| LAI_InB | 7.04 (1.22) | 5.97 | 8.77 | 1.73 | 18.83 | * | * | ms |
| LAI_A7 | 5.35 (0.84) | 4.38 | 6.23 | 1.22 | 19.05 | ms | ns | ns |
| k_InB | 0.46 (0.09) | 0.38 | 0.54 | 0.13 | 19.95 | ns | * | * |
| k_A7 | 0.78 (0.15) | 0.67 | 0.91 | 0.16 | 16.66 | * | ms | ns |
| InB (days) | 61 (0.96) | 58 | 65 | 2.17 | 1.84 | *** | *** | *** |
| H (days) | 71 (0.92) | 68 | 75 | 2.37 | 1.31 | *** | *** | *** |
| A7 (days) | 76 (1.15) | 73 | 80 | 2.52 | 1.61 | *** | *** | *** |
| PM (days) | 116 (1.01) | 113 | 120 | 2.4 | 1.01 | *** | *** | *** |

HI: harvest index, TGW: thousand grain weight, GSP: grains per spike, GWSP: grain weight per spike, GM2: grains per $m^{-2}$, SM2: spikes per $m^{-2}$, LI: light interception, LAI: leaf area index, k: extinction coefficient, RUE_preGF: radiation use efficiency from pre grain filling period, RUE_GF: radiation use efficiency from grain filling period, RUE_Total: radiation use efficiency from the whole crop cycle, BM: aboveground biomass, WSC: water soluble carbohydrates.

Supplemental Table 2. Statistical significance of the main effects genotype (G), year (Y), growth stage (GS) and leaf position within the canopy (P) and their interactions on the physiological traits measured in this study. * = significant at $p<0.05$, ** = significant at $p<0.01$, *** = significant at $p<0.001$, ms = marginally significant ($0.1>p>0.05$), ns = no statistical significance.

| Trait | G | Y | GS | P | GxY | GxGS | YxGS | GxP | YxP | GSxP | GxYxGS | GxYxP | GxGSxP | YxGSxP | GxYxGSxP |
|---|---|---|---|---|---|---|---|---|---|---|---|---|---|---|---|
| $A_{sat}$ | *** | *** | *** | *** | *** | ms | ns | ns | * | *** | ms | ns | ns | ns | ns |
| LAI | *** | *** | *** | *** | *** | *** | *** | ms | *** | *** | ** | ns | ns | *** | ns |
| Length | *** | *** | *** | *** | *** | * | *** | *** | * | ns | * | *** | ns | ns | ns |
| Width | *** | *** | *** | *** | *** | * | *** | *** | *** | ns | *** | ** | ns | ns | ns |
| SLA | *** | *** | *** | *** | * | *** | *** | ** | *** | *** | * | ** | *** | *** | ** |
| LI | ns | *** | *** | *** | ns | ns | ms | *** | *** | *** | ns | *** | *** | *** | * |
| SPAD | *** | * | *** | *** | * | ** | ** | * | *** | *** | ns | ns | ns | *** | ns |
| N content | ns | *** | *** | *** | ns | ** | ns | ns | ns | ** | *** | ns | ns | *** | ns |
| C content | ns | ns | *** | ns | ms | ns | ns | ns | * | * | ** | ns | ns | * | ns |
| SLN | *** | *** | *** | *** | ns | ** | ** | ** | *** | *** | ** | ns | ns | *** | ns |
| C:N | *** | *** | *** | *** | ns | ** | ms | ns | ns | *** | ** | ns | ns | ** | ns |

$A_{sat}$: $CO_2$ assimilation at light saturating conditions, LAI: Leaf Area Index, SLA: Specific Leaf Area, LI: %Light Interception, SLN: Specific Leaf Nitrogen, C:N: Carbon nitrogen ratio.

Supplemental Table 3. Relationships between gas exchange traits ($CO_2$ assimilation under light saturating conditions [$A_{sat}$], stomatal conductance [*gs*] and dark respiration [$R_d$]) with yield, and aboveground biomass in staple crops. YP: Yield potential, NT: N treatments, D: Drought. Where the study did not specifically look for correlations between $A_{sat}$, *gs* or $R_d$ with biomass and/or yield the tool WebPlotDigitizer (http://arohatgi.info/WebPlotDigitizer) was used to extract the values and calculate the regression coefficient between the variables. When not specified, the photosynthetic measurements were done in the uppermost leaf at the specific growth stage measurement took place.

| Trait | Yield | Biomass at maturity | Crop | Environment | Reference |
|---|---|---|---|---|---|
| $A_{sat}$ | $r^2$ = 0.24 (top layer booting)<br>$r^2$ = 0.11 (middle layer booting)<br>$r^2$ = 0.33 (bottom layer booting)<br>$r^2$ = 0.54, $p<0.01$ (top layer grain filling)<br>$r^2$ = 0.05 (middle layer grain filling)<br>$r^2$ = 0.23 (bottom layer grain filling)<br>$r^2$ = 0.12 (canopy booting)<br>$r^2$ = 0.13 (canopy grain filling) | $r^2$ = 0.01 (top layer booting)<br>$r^2$ = 0.02 (middle layer booting)<br>$r^2$ = 0 (bottom layer booting)<br>$r^2$ = 0.06 (top layer grain filling)<br>$r^2$ = 0.09 (middle layer grain filling)<br>$r^2$ = 0.02 (bottom layer grain filling)<br>$r^2$ = 0.13 (canopy booting)<br>$r^2$ = 0.03 (canopy grain filling) | Wheat | YP | This study |
| | $r^2$ = 0.4, $p<0.01$ (4 months after planting) | $r^2$ = 0.37, $p<0.01$ (4 months after planting) | Cassava | YP | (El-Sharkawy et al., 1990) |
| | $r^2$ = 0.35, $p<0.01$ (panicle initiation YP)<br>$r^2$ = 0.86, $p<0.01$ (mid-development YP)<br>$r^2$ = 0.53, $p<0.01$ (head exertion YP)<br>$r^2$ = 0.74, $p<0.001$ (avg YP)<br>$r^2$ = 0.25, $p<0.05$ (panicle initiation D)<br>$r^2$ = 0.44, $p<0.01$ (mid-development D)<br>$r^2$ = 0.13 (head exertion D)<br>$r^2$ = 0.5, $p<0.001$ (avg D) | $r^2$ = 0.32, $p<0.01$ (panicle initiation YP)<br>$r^2$ = 0.83, $p<0.01$ (mid-development YP)<br>$r^2$ = 0.46, $p<0.01$ (head exertion YP)<br>$r^2$ = 0.67, $p<0.001$ (avg YP)<br>$r^2$ = 0.36, $p<0.05$ (panicle initiation D)<br>$r^2$ = 0.55, $p<0.01$ (mid-development D)<br>$r^2$ = 0.15 (head exertion D)<br>$r^2$ = 0.66, $p<0.001$ (avg D) | Sorghum | YP and D | (Peng et al., 1991) |
| | $r^2$ = 0 (YP)<br>$r^2$ = 0.4, $p<0.01$ (D) | $r^2$ = 0.01 (YP)<br>$r^2$ = 0.53, $p<0.01$ (D) | Wheat | YP and D | (Wada et al., 1994) |
| | $r^2$ = 0.13 (pre-anthesis)<br>$r^2$ = 0.84, $p<0.001$ (post-anthesis) | $r^2$ = 0.25 (pre-anthesis)<br>$r^2$ = 0.74, $p<0.001$ (post-anthesis) | Wheat | YP | (Gent, 1995) |
| | $r^2$ = 0.85, $p<0.05$ (average pre and post-anthesis) | $r^2$ = 0.07 (average pre and post-anthesis) | Wheat | YP | (Fischer et al., 1998) |
| | $r^2$ =0.28, $p<0.05$ (post-anthesis) | $r^2$ = 0.64, $p<0.01$ (post-anthesis) | Wheat | YP | (Gutiérrez-Rodríguez et al., 2000) |

| | | | | |
|---|---|---|---|---|
| $r^2$ =0.52, $p<0.01$ (average pre and post-anthesis) | $r^2$ = 0.36, $p<0.01$ (average pre and post-anthesis) | Wheat | YP | (Reynolds et al., 2000) |
| $r^2$ = 0.52, $p<0.001$ (mid-stem elongation)<br>$r^2$ = 0.31, $p<0.05$ (late stem elongation)<br>$r^2$ = 0.19, $p<0.1$ (heading)<br>$r^2$ = 0.44, $p<0.01$ (anthesis)<br>$r^2$ = 0.28, $p<0.05$ (soft dough)<br>$r^2$ = 0.04 (hard dough)<br>$r^2$ = 0.3, $p<0.05$ (cycle average) | $r^2$ = 0.03 (mid-stem elongation)<br>$r^2$ = 0.01 (late stem elongation)<br>$r^2$ = 0.13 (heading)<br>$r^2$ = 0.01 (anthesis)<br>$r^2$ = 0 (soft dough)<br>$r^2$ = 0.01 (hard dough)<br>$r^2$ = 0.03 (cycle average) | Wheat | YP | (Jiang et al., 2003) |
| | $r^2$ = 0.04 | Wheat | YP | (Chytyk et al., 2011) |
| $r^2$ = 0.42, $p<0.01$ (grain filling) | | Wheat | YP | (Zheng et al., 2011) |
| $r^2$ = 0.12 | $r^2$ = 0.07 | Wheat | YP | (Driever et al., 2014) |
| $r^2$ = 0.56, $p<0.05$ (heading, YP)<br>$r^2$ = 0.01 (3 days after anthesis, YP)<br>$r^2$ = 0.26 (20 days after anthesis, YP)<br>$r^2$ = 0.49 (heading, D)<br>$r^2$ = 0 (3 days after anthesis, D)<br>$r^2$ = 0.12 (20 days after anthesis, D) | | Wheat | YP and D | (Sun et al., 2014) |
| $r^2$ = 0.49 (jointing)<br>$r^2$ = 0.01 (anthesis)<br>$r^2$ = 0.34 (grain filling)<br>$r^2$ = 0.23 (average) | $r^2$ = 0.02 (jointing)<br>$r^2$ = 0.15 (anthesis)<br>$r^2$ = 0.05 (grain filling)<br>$r^2$ = 0.07 (average) | Wheat | YP | (Chen and Hao, 2015) |
| $r^2$ = 0.98 (average)<br>$r^2$ = 0.93 (moderate N)<br>$r^2$ = 0.98 (high N) | $r^2$ = 0.89 (average)<br>$r^2$ = 0.86 (moderate N)<br>$r^2$ = 0.92 (high N) | Rice | DNT | (Huang et al., 2016) |
| $r^2$ = 0.75, $p<0.001$ (pre-anthesis)<br>$r^2$ = 0.76, $p<0.001$ (post-anthesis) | $r^2$ = 0.63, $p<0.001$ (pre-anthesis)<br>$r^2$ = 0.59, $p<0.001$ (post-anthesis) | Wheat | DNT | (Gaju et al., 2016) |
| $r^2$ = 0 ($A_{sat}$)<br>$r^2$ = 0.27, $p<0.05$ (AQ1000, pre-anthesis)<br>$r^2$ = 0.27, $p<0.05$ (AQ1000, post-anthesis) | $r^2$ = 0 | Wheat | YP | (Carmo-Silva et al., 2017) |
| $r^2$ = 0.9, $p<0.01$ | | Rice | YP | (Chen et al., 2020) |
| $r^2$ = 0.12 (spike)<br>$r^2$ = 0.17 (flag leaf) | | Wheat | YP | (Molero and Reynolds, 2020) |

| | | | | | |
|---|---|---|---|---|---|
| *gs* | $r^2$ = 0.04 (top layer booting)<br>$r^2$ = 0.05 (middle layer booting)<br>$r^2$ = 0 (bottom layer booting)<br>$r^2$ = 0.01 (top layer grain filling)<br>$r^2$ = 0 (middle layer grain filling)<br>$r^2$ = 0.07 (bottom layer grain filling)<br>$r^2$ = 0.04 (canopy booting)<br>$r^2$ = 0.01 (canopy grain filling) | $r^2$ = 0.05 (top layer booting)<br>$r^2$ = 0.14 (middle layer booting)<br>$r^2$ = 0.07 (bottom layer booting)<br>$r^2$ = 0.21 (top layer grain filling)<br>$r^2$ = 0.52, $p<0.01$ (middle layer grain filling)<br>$r^2$ = 0.21 (bottom layer grain filling)<br>$r^2$ = 0.03 (canopy booting)<br>$r^2$ = 0.34, $p<0.1$ (canopy grain filling) | Wheat | YP | This study |
| | $r^2$ = 0.03 | $r^2$ = 0.01 | Cassava | YP | (El-Sharkawy et al., 1990) |
| | $r^2$ = 0.85, $p<0.001$ (average pre and post-anthesis) | $r^2$ = 0.01 (average pre and post-anthesis) | Wheat | YP | (Fischer et al., 1998) |
| | $r^2$ =0.28, $p<0.05$ (post-anthesis) | $r^2$ = 0.58, $p<0.01$ (post-anthesis) | Wheat | YP | (Gutiérrez-Rodríguez et al., 2000) |
| | $r^2$ = 0.72, $p<0.01$ (average pre and post-anthesis) | $r^2$ = 0.45, $p<0.01$ (average pre and post-anthesis) | Wheat | YP | (Reynolds et al., 2000) |
| | $r^2$ = 0.53, $p<0.001$ (mid-stem elongation)<br>$r^2$ = 0.24, $p<0.05$ (late stem elongation)<br>$r^2$ = 0.27, $p<0.05$ (heading)<br>$r^2$ = 0.26, $p<0.05$ (anthesis)<br>$r^2$ = 0.21 (soft dough)<br>$r^2$ = 0.12 (hard dough)<br>$r^2$ = 0.27, $p<0.05$ (cycle average) | $r^2$ = 0.09 (mid-stem elongation)<br>$r^2$ = 0.24, $p<0.05$ (late stem elongation)<br>$r^2$ = 0.01 (heading)<br>$r^2$ = 0.07 (anthesis)<br>$r^2$ = 0 (soft dough)<br>$r^2$ = 0.01 (hard dough)<br>$r^2$ = 0.07 (cycle average) | Wheat | YP | (Jiang et al., 2003) |
| | | $r^2$ = 0.25 (jointing)<br>$r^2$ = 0 (flowering)<br>$r^2$ = 0.01 (grain filling)<br>$r^2$ = 0.08 (average) | Wheat | YP | (Chytyk et al., 2011) |
| | $r^2$ = 0.48, $p<0.01$ | | Wheat | YP | (Zheng et al., 2011) |
| | | $r^2$ = 0.78 | Wheat | YP | (Pang et al., 2014) |
| | $r^2$ = 0.77, $p<0.05$ (jointing)<br>$r^2$ = 0.08 (anthesis)<br>$r^2$ = 0.04 (grain filling)<br>$r^2$ = 0 (average) | $r^2$ = 0.25 (jointing)<br>$r^2$ = 0 (anthesis)<br>$r^2$ = 0.01 (grain filling)<br>$r^2$ = 0.08 (average) | Wheat | YP | (Chen and Hao, 2015) |

| | | | | | |
|---|---|---|---|---|---|
| | $r^2 = 0.39$, $p<0.05$ (pre-anthesis)<br>$r^2 = 0.37$, $p<0.05$ (post-anthesis) | $r^2 = 0.39$, $p<0.05$ (pre-anthesis)<br>$r^2 = 0.34$, $p<0.05$ (post-anthesis) | Wheat | DNT | (Gaju et al., 2016) |
| | $r^2 = 0$ | $r^2 = 0$ | Wheat | YP | (Carmo-Silva et al., 2017) |
| $R_d$ | $r^2 = 0$ (average pre and post-anthesis) | $r^2 = 0.02$ (average pre and post-anthesis) | Wheat | YP | (Reynolds et al., 2000) |
| | | $r^2 = 0.25$ | Wheat | YP | (Chytyk et al., 2011) |
| | $r^2 = 0.05$ (spike, heading)<br>$r^2= 0.46$ (spike, grain filling)<br>$r^2 = 0.26$ (spike, average) | $r^2 = 0$ (spike, heading)<br>$r^2= 0.22$ (spike, grain filling)<br>$r^2 = 0.11$ (spike, average) | Wheat | YP | (Zhou et al., 2016) |
